# Patch deconvolution for Fourier light-field microscopy

**DOI:** 10.1101/2025.06.13.659385

**Authors:** Bin Fu, Caroline L. Jones, Daniel Heraghty, Shengbo Yang, Caitlin O’Brien-Ball, Victoria Junghans, Haowei Yang, Tuomas P.J. Knowles, Lucien E. Weiss, Ricardo A. Fernandes, Steven F. Lee

## Abstract

Imaging flow cytometry using Fourier light-field microscopy enables high-throughput three-dimensional cellular imaging, capable of capturing thousands of events per second. However, volumetric reconstruction speed remains orders of magnitude slower than the acquisition speed. The current state of art uses Richardson–Lucy algorithm, restricted to just 5-10 reconstructed per second with GPU acceleration. This limitation hinders real-time applications such as cell sorting and thus has bottlenecked the widespread adoption of 3D imaging flow cytometry. We introduce patch deconvolution, an optimisation compatible with the Richardson–Lucy framework that significantly accelerates convergence, achieving over 100-200 reconstructions per second on standard GPUs, a 20–40 fold improvement over Richardson–Lucy. Validated on both simulated and experimental datasets, patch deconvolution achieves reconstruction quality comparable to Richardson–Lucy in both static and flow data. This supports rapid cell sorting based on spatial features and enables advanced applications, such as detecting rare spatial events in large cell populations, which would otherwise be indistinguishable in traditional flow cytometry.

## 1. Introduction

Over the past 60 years, flow cytometry has become an indispensable tool for cell biologists to interrogate population-scale data [1]. By measuring the light scattered and emitted as cells pass through a focused light beam, flow cytometers along with additional hardware and control systems can also sort cells quickly. For instance, fluorescence-activated cell sorting (FACS) enables high-throughput sorting of thousands of cells per second but is limited to low-resolution parameter spaces, *e.g*.intensity and size [1]. Alternatively, fluorescence microscopy enables detailed spatial information but generally has low throughput. Imaging flow cytometry (IFC) combines the strengths of both techniques, merging the speed of flow cytometry with the spatial detail of microscopy [2–4]. As such, IFC allows investigation of population-level dynamics based on complex spatial metrics of individual cells such as eccentricity, spatial clustering and correlation between fluorescent labels [5, 6]. IFC, such as developed by Schraivogel *et al*. [7]. achieves cell sorting at rates of 15000 events/s using a low-latency image processing algorithm to make real-time, image-based sorting decisions. However, IFC approaches rely on 2D images of cells, limiting their ability to capture the full 3D spatial information, which is often critical for understanding cellular structure and function in biological systems [3], such as protein traffics [8], protein-protein colocalisations [9].

Efforts to extend flow cytometry to high-throughput 3D imaging have shown promise but face significant trade-offs, including reduced throughput (<100 events per second) [10, 11], increased system complexity [12–14], or limited depth of field and emitter density [15]. Light-field flow cytometry (LFC), the implementation using Fourier light-field microscopy (FLFM) [16–19] as demonstrated by Hua and Han *et al*., addresses these challenges. FLFM captures multiple perspectives of the sample in a single snapshot, achieving throughputs of up to 5000 events/s on a 512×512 image [20]. However, accurate reconstruction relies heavily on computationally intensive deconvolution algorithms. Richardson–Lucy (RL) deconvolution [21, 22], an iterative expectation–maximization (EM) algorithm [23, 24], is widely used due to its robustness and simplicity [25]. In the system described by Hua and Han *et al*., a 4 µm depth of field (DOF) requires approximately 10 seconds per volume for RL deconvolution. To capture entire cells, a greater number of microlenses is required for a larger DOF, resulting in a larger image and volume size and a corresponding quadratic increase in reconstruction time. This creates a bottleneck, where reconstruction times, often taking days for large datasets, far exceed imaging times, thus limiting LFC’s practical use in real-time applications such as cell sorting.

Various strategies have been explored to accelerate RL deconvolution, including algorithmic optimisations, deep learning approaches, and hardware-efficient matrix optimisations. Algorithmic methods focus on accelerating convergence by modifying the objective function [26–29], such as incorporating Wiener–Butterworth filters [28], which can reduce the number of iterations to fewer than 5. Nonetheless, even a single iteration typically requires tens of milliseconds and requires a large GPU memory, making it impractical for high-throughput applications and also it was proved to be less effective in FLFM reconstruction [29]. Deep learning-based reconstruction methods [30–33] also have own limitations. Aside from the challenge of obtaining large, high-quality ground truth datasets for training, these models often exhibit poor generalisability across varying sample structures, noise levels, and optical aberrations as demonstrated by Lu *et al*. [33]. Although SeReNet [33], a self-supervised architecture, demonstrates improved generalisability, it still requires hundreds of milliseconds per volume reconstruction. Additionally, matrix-level optimisations [34–36], such as reformulating the RL update step to use more efficient matrix structures [36], offer gains in speed and memory efficiency, particularly on GPU architectures, yet reconstruction times still require tens of iterations to converge and thus remain in the range of hundreds of milliseconds.

To further reduce reconstruction time to the millisecond scale while preserving the advantages of existing optimisation strategies, we introduce patch deconvolution, a technique fully compatible with conventional RL deconvolution frameworks and significantly improving computational efficiency without compromising reconstruction accuracy. Inspired from Bayesian-based multiview deconvolution in light-sheet microscopy [37] and ordered-subsets EM (OSEM) in computed tomography [38], patch deconvolution reformulates the update process by sequentially using individual perspective views, rather than processing the entire FLFM image at once, during each iteration. This strategy enables multiple volume updates per PSF, proportional to the number of views captured in the FLFM system. In contrast, standard RL deconvolution performs a single update per iteration using the full PSF. Consequently, patch deconvolution achieves a speedup factor theoretically equal to the number of views, which at least has to be 19 for a whole cell imaging (Supplementary Note S1.A), allowing complete 3D reconstructions to be performed in milliseconds on a GPU. Furthermore, because the method is grounded in the RL framework, it remains fully compatible with existing algorithm-level and matrix-level optimisations and also regularisations [39, 40].

In this work, we describe the principle of patch deconvolution and its implementation within existing RL framework. We validate its performance through simulations and experimental datasets (both static and in-flow FLFM at acquisition rates of 300–1500 events/s). Our results demonstrate strong agreement between patch deconvolution and conventional RL reconstructions, both qualitatively and quantitatively. Furthermore, we provide a mathematical proof that, under noise-free and aberration-free conditions, patch deconvolution converges to the same ground truth solution as standard RL deconvolution (Supplementary Note S2.D). Critically, patch deconvolution enables, for the first time, 3D reconstruction in a millisecond scale for LFC, thereby supporting the highest cell-sorting throughput currently achievable based on 3D data.

## 2. Principle of patch deconvolution

RL deconvolution, derived from Bayes’ theorem and implemented as an iterative expectation-maximisation (EM) algorithm, has been widely employed in astronomy and medical imaging for image restoration due to its simplicity and efficacy [40]. Given the point-spread function (PSF) of a system, *h* (*s*), and the observed image, *I* (*s*), where *s* denotes each pixel, RL deconvolution iteratively estimates the latent image by maximising the likelihood of the estimated data based on the measured data. Each iteration comprises four sequential steps: forward propagation (FP); convolving the current estimate, *o*_*k*_ (*s*), with *k* representing the iteration, with the PSF, *h* (*s*), to simulate the measured image. Error comparison (EC); the ratio between the measured image, *I* (*s*), and the FP result. Back propagation (BP); convolving the EC result with the flipped PSF *h* (−*s*) and the result update (RU); multiplying the BP result with the current estimate for updating (Fig. 2a). For implementation of RL deconvolution in FLFM (Fig. 2b), the BP step is represented in Eqn. 1 and RU step in Eqn. 2, where *f*_RL_ (*s*; *z*) denotes the back-projected image at a specific axial plane, and the integral characterises the projection of the 3D volume into a 2D image [41].

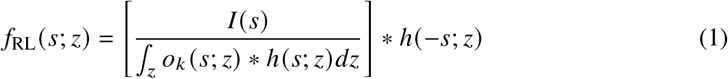

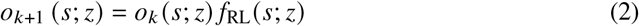

**Fig. 1.**
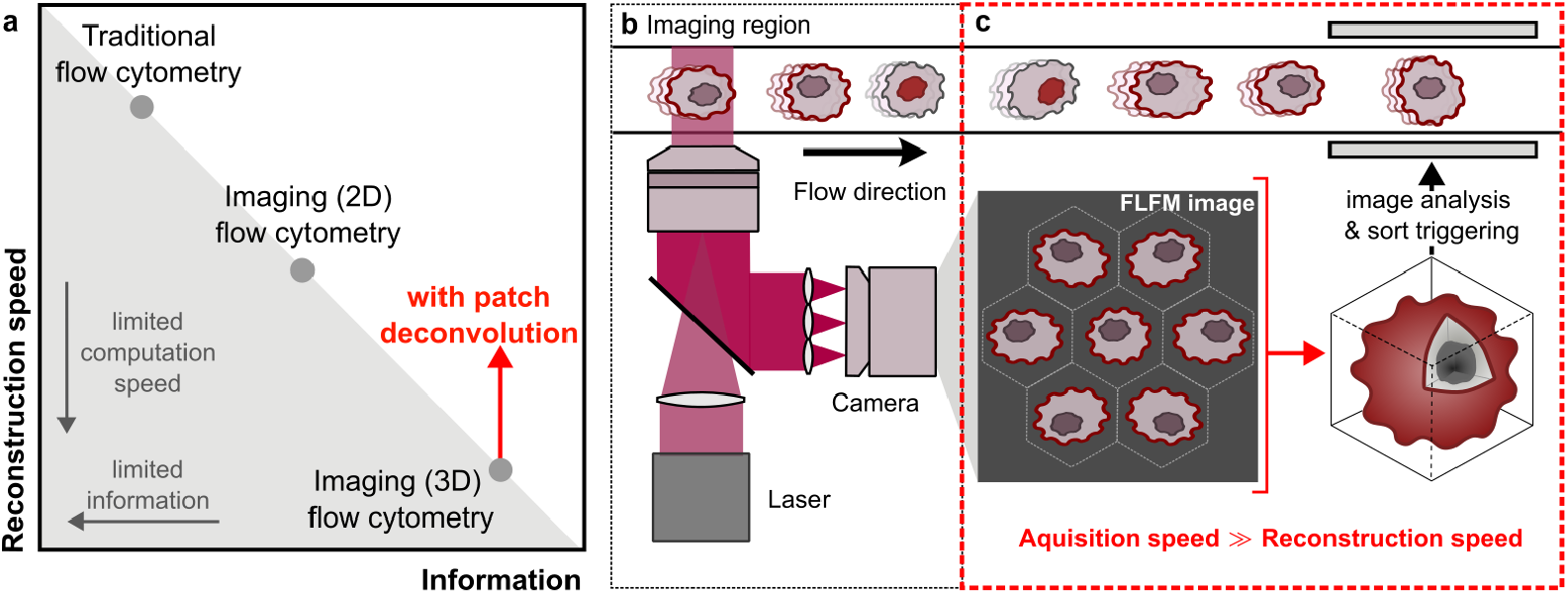
Patch deconvolution enables cell sorting for 3D imaging flow cytometry. **a** Comparison of traditional flow cytometry, 2D imaging flow cytometry, and 3D imaging flow cytometry in terms of reconstruction speed and information content. The bottleneck in 3D imaging cytometry arises from the slow reconstruction speeds. Patch deconvolution (red arrow) is proposed to address this, enabling faster reconstruction speed for 3D flow cytometry. **b** Schematic of a 3D flow cytometry setup using a high NA objective, laser illumination, and a microlens array (MLA) to capture a FLFM image for flowing cells. **c** Workflow of FLFM image acquisition and image reconstruction for sorting. Patch deconvolution accelerates the time-intensive image reconstruction step.

**Fig. 2.**
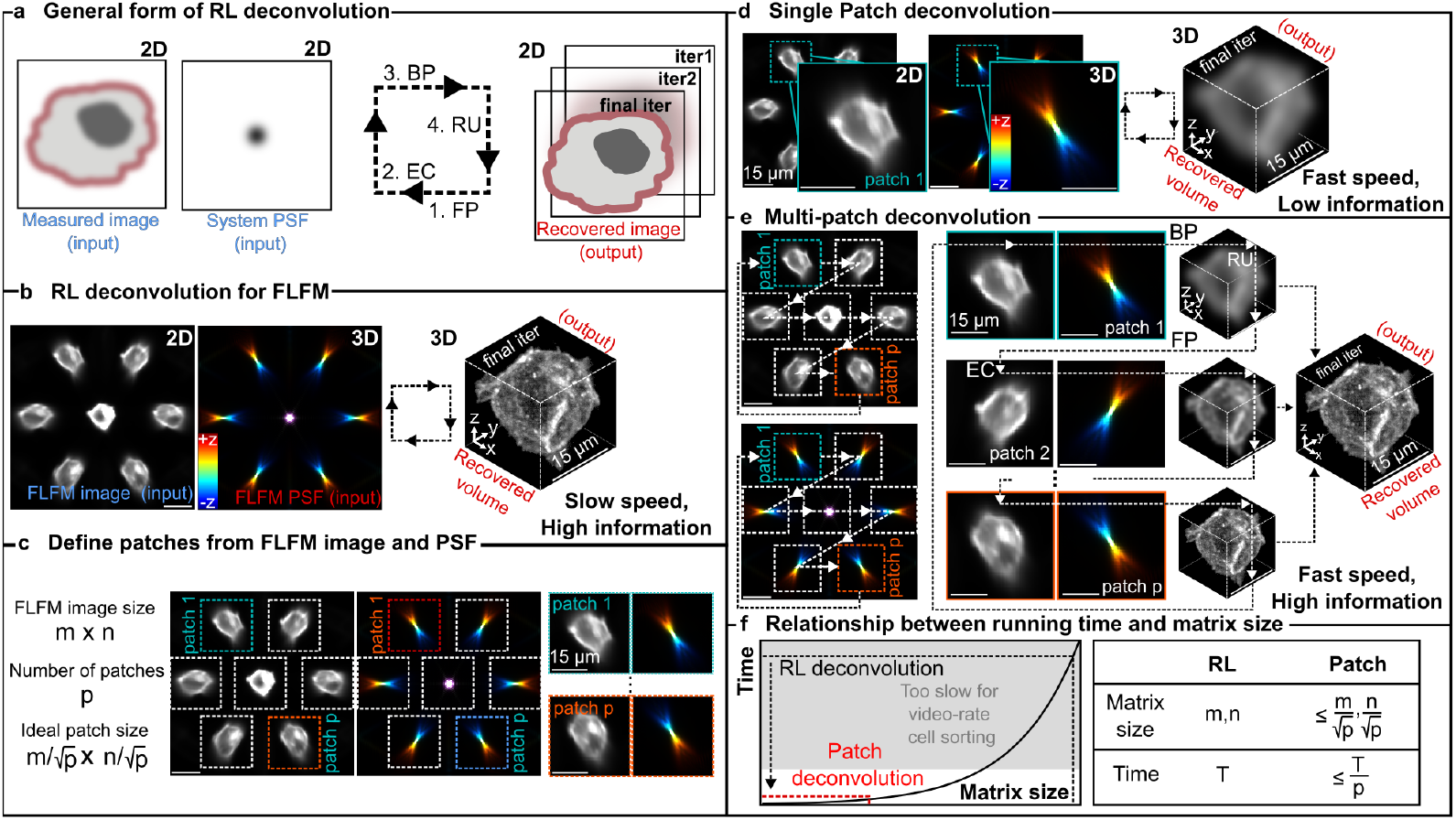
RL deconvolution and patch deconvolution for FLFM reconstruction. **a** RL deconvolution involves iterative forward projection (FP), error correction (EC), back projection (BP), and result update (RU). **b** In FLFM, 3D volumes are reconstructed iteratively from the measured image and PSF but are computationally slow. **c** Dividing images and PSFs into patches reduces computation; optimal patch size is 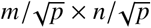. **d** Single-patch deconvolution is fast but with low result quality. **e** Multi-patch deconvolution (also referred to as patched deconvolution in this paper) method. The FLFM image and PSF are divided into multiple patches, with deconvolution performed independently on each patch in sequence. **f** Reconstruction runtime scales quadratically with matrix size. Therefore, patching can significantly reduce time.

Building on the RL deconvolution framework, we introduce the patch deconvolution that takes the advantage of the multi-view nature of FLFM, similar to approaches in multi-view light-sheet microscopy and computed tomography. In FLFM, a single image comprises multiple perspectives of the same object, resulting from the MLA located in the BFP. Each of these perspective views, or *patches*, is effectively independent and associated with its own unique PSF. Consequently, each patch can be deconvolved individually. If an FLFM image contains *p* perspectives, it can be partitioned into *p* patches, each ideally of size 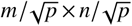, where (*m* × *n*)denotes the dimensions of the original image (Fig. 2c).

In patch deconvolution (Fig. 2e), the BP step from a single patch is described in Eqn. 3, and the RU step sequentially using views to update reconstructed volume is formalised in Eqn. 4. Here, one iteration is defined as processing one patch, with *P* representing the full set of patches and *p* a specific patch. A mathematical convergence prove is in Supplementary Note S2.D. For implementation in FLFM, we adopt a plane-wise optimisation approach [36]. Instead of performing a full 3D deconvolution by padding the 2D FLFM data into a 3D volume, we apply 2D deconvolution using a 2D PSF at each axial plane. This approach enables recovery of 2D volume slices corresponding to each plane, significantly reducing the memory requirements on the GPU.

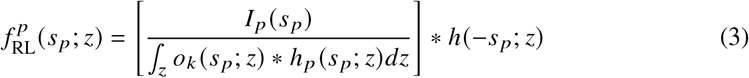

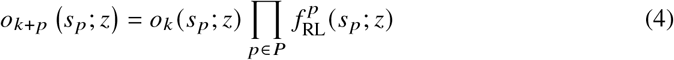

## 3. Materials and Methods

### 3.1. Experimental setup

Experiments were conducted using one of two distinct instruments: a wide-field fluorescence microscope with a MLA for FLFM imaging and a spinning-disk confocal microscope detailed in [42] for acquiring high-resolution data for simulation.

The FLFM microscope was a bespoke widefield fluorescence microscope, with the illumination entering the microscope body through the back illumination port. The excitation path has a 638nm laser (Odic force lasers, DM-RL500). The laser beam was circularly polarized using quarter-wave plates. Light is then focused onto the back focal plane of an oil immersion objective (Olympus, UPlanSApo 60x/1.30 NA silicon) via a dichroic mirror (Semrock, Di01-R405/488/561/635). Fluorescence emitted from the sample is filtered by an emission filter (Semrock, FF01-432/515/595/730) before passing through a 100mm or 125 mm Fourier lens (Thorlabs AC254-100-A-ML and AC254-125-A-ML for 19 and 37 MLA configurations, respectively), positioned at a focal length from the native image plane. A MLA, pitch 1 mm and focal length 36.7mm (Okotech, APH-Q-P1000-F36,7), is placed at a focal length from the Fourier lens. A high-speed sCMOS camera (Photometrics, Kinetix) is positioned at the MLA image plane.

A custom flow-focusing microfluidic device were used in this work. It was fabricated from PDMS on glass coverslips using established protocols [43]. An ElveFlow OB1 microfluidic flow controller was used for individual control of the separate sheath and sample channels. A Thorlabs air compressor (Thorlabs, PTA522) was used to pressurise the system. Sheath fluid (PBS) and sample were kept in sealed eppendorfs and delivered independently to the microfluidic device by 1.6 mm OD PTFE tubing (BL-PTFE-1608-20, Darwin Microfluidics) wiht a 20mbar pressure. Cells were resuspended in 1ml of PBS, which was plumbed into the microfluidic system alongside a separate 1ml Eppendorf of PBS for the sheath channel.

### 3.2. Light-field image formation

The 3D PSF of the FLFM microscope was simulated using formulas described in Supplementary Note S2.A. The PSF voxel size was set to be the same as the camera pixel size at the object plane across a 20 or 30 µm axial range for the 19 MLA and 37 MLA configurations, respectively. To ensure a full sampling of PSF, the simulation was performed over a matrix larger than the BFP, using a size of 1000 × 1000 pixels.

The ground truth objects were 3D cell volumes cropped and resampled from confocal microscopy data. These volumes were zero-padded in all three dimensions to match the simulated PSF matrix dimensions. The simulated FLFM image was then generated using Eqn. 5, where *I* denotes the simulated FLFM image. These images were normalised to a range between 0 and 1. FLFM images with different levels of noise were then created by adding a series of Poisson-noise images with varying standard deviation (*σ*), to the original noiseless image. The peak signal-to-noise ratio (PSNR) of the resulting FLFM images were calculated as 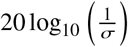.

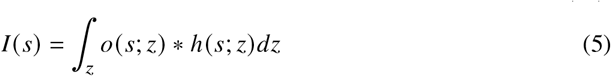

### 3.3. Reconstruction for static and flow data

The reconstruction was performed using a self-implemented version of RL deconvolution and patch deconvolution. The number of iterations in the RL deconvolution was set equal to the number of patches used in the patch deconvolution, which was equal to the number of views.

All reconstructions were performed using MATLAB 2024a on a PC with an Intel i7-13700K CPU and Nvidia RTX 4060 GPU. For RL deconvolution, the reconstructed volume had the same matrix dimensions as the simulated PSF. In the case of patch deconvolution, the axial dimension of both the volume and PSF matched that used in RL deconvolution. However, the lateral dimensions were limited to 130×130 pixels, corresponding to field-of-view (FOV) size in pixel. After reconstruction, the volume obtained from the RL deconvolution was cropped from the center to match the size of the volume reconstructed from the patch deconvolution. Both RL and patch volumes were normalized between 0 and 1. For experimental datasets, a hybrid PSF (Supplementary Note S5) was used during the reconstruction process.

### 3.4. Sample preparation

Jurkat cells were used in this work for performance comparison. Membrane receptor CD45 on the surface of Jurkat cells was labelled with antibody gap 8.3 and AlexaFluor 647 (concentration 15 µg/ml). Cells were cultured in RPMI-1640 media (Merck/Sigma-Aldrich Cat No: R2405-500ML) with 10% FBS (ThermoFisher REF:10500-064), 1% sodium pyruvate (ThermoFisher REF:11360-039), 1% HEPES (ThermoFisher REF:15630-056) and 1% penicillin-streptomycin (ThermoFisher REF: 15140122). Following three washes in 20 nm filtered PBS cells were suspended in 30 µl of the antibody solution and incubated for 15 minutes. Cells were then washed twice more in PBS before fixation in 0.8% PFA (thermo scientific, 28906) and 0.2% glutaraldehyde (Sigma-Aldrich, 3802-75ML) solution. They were left to fix at room temperature for 15 minutes. Cells were washed two final times before resuspension in PBS for imaging. Cells were plated in a µ-Slide 18 Well Glass Bottom slide (ibidi, 80807).

## 4. Result

### 4.1. Validation on static bead and cell data

To evaluate the performance of patch deconvolution, we first tested it on simulated data without noise and aberrations (Fig. 3a and Supplementary Note S3). Under these conditions, patch deconvolution converged to a result nearly identical to that of RL deconvolution when using the number of iterations equal to the number of views in the FLFM image (*i.e*.each patch was used once).

**Fig. 3.**
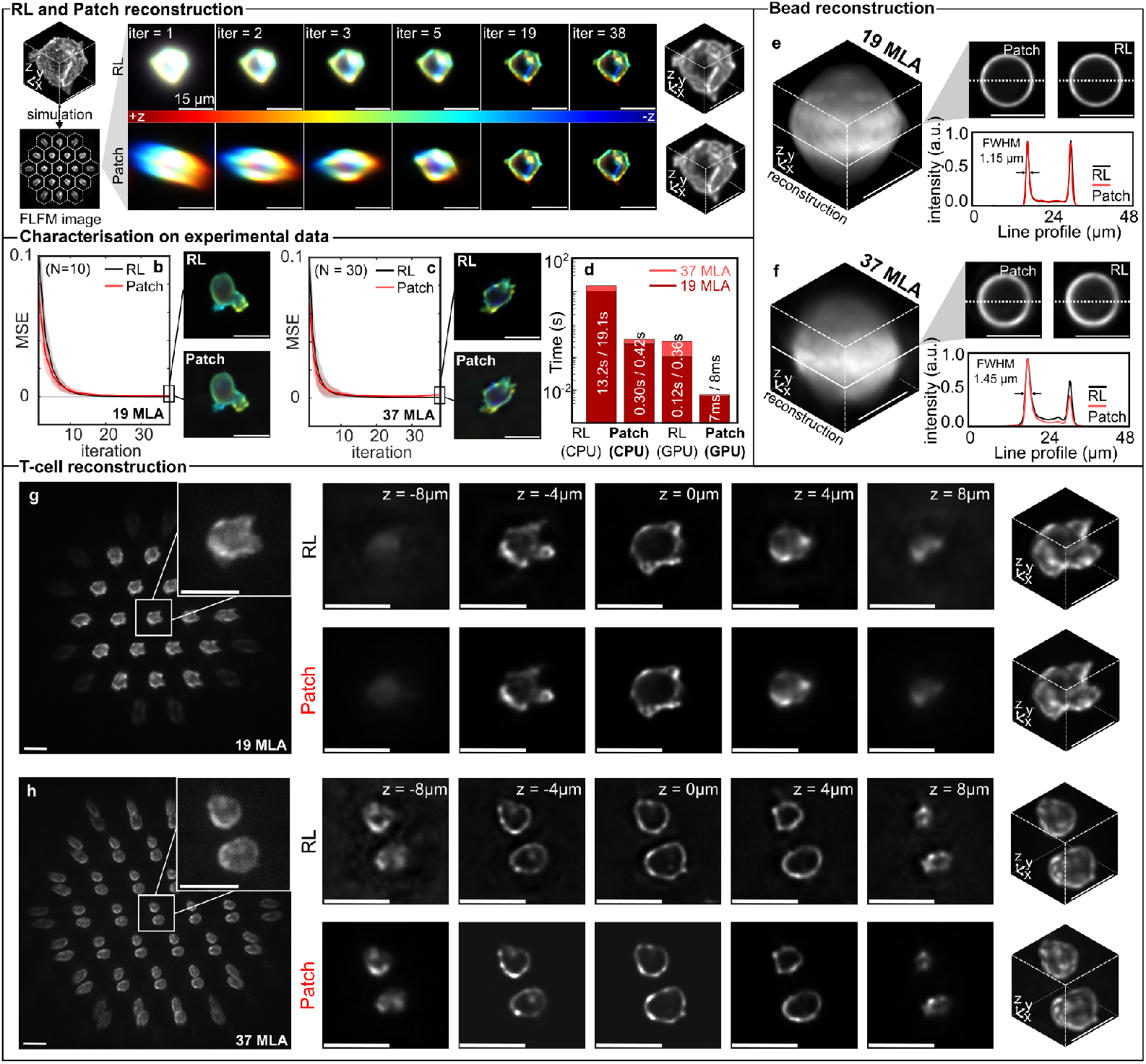
Validation on static data. **a** Comparison of RL (top row) and patch (bottom row) deconvolution on the same simulated FLFM image with 19 sub-aperture views. **b, c** MSE between RL and patch deconvolution for experimental T-cell data using 19 and 37 MLA configurations; RL results at the 19^th^ and 37^th^ iterations were used as reference. **d** Runtime comparison of both methods over 37 iterations on CPU and GPU for 19 and 37 MLA cases. **e, f** Bead reconstruction comparison (volume, 2D slice, and line profile) between RL and patch deconvolution for a 15 µm bead. **g, h** T-cell reconstructions with 19 and 37 MLA show consistent results between methods across the full cell depth. All scale bars are 15 µm.

We next validated the patch deconvolution on experimental data, imaging static beads and membrane-labelled T-cells using the FLFM microscope with a 100 ms exposure. For bead imaging, fluorescently surface-labelled 15 µm beads (Thermofisher, FocalCheck™ Fluorescence Microscope Test Slide #1) were imaged in both 19 MLA and 37 MLA configurations. Reconstructions were performed using a hybrid PSF that accounted for system aberrations (Supplementary Note S5), with the same number of iterations for both patch and RL deconvolution, equal to the number of views. Figures 3e and f show the 3D reconstructions after 19 and 37 iterations, accurately reflecting bead size and shape. The 2D *xy*-slices at *z* = 0 exhibit similar line profiles and identical full-width half-maximum (FWHM) for both methods. A library of bead reconstructions is presented in Supplementary Note S4.A.

T-cell data were acquired using highly inclined and laminated optical sheet illumination (HiLO) in both MLA configurations. Again, the number of iterations matched the number of views. The final RL deconvolution result was used as the reference for mean squared error (MSE) calculation. Convergence based on MSE are shown in Figs. 3b and c, indicating that patch deconvolution reached results comparable to RL deconvolution in high signal conditions (19 MLA: 8.4 ×10^−4^; 37 MLA: 1.8 ×10^−3^). Pixel-wise convergence across depth is shown in Fig. 3g and h, demonstrating that patch deconvolution accurately reconstructed cell morphology across the field of view within the axial range of a whole cell, albeit with slightly elevated background noise. Runtime benchmarking was performed on an Nvidia RTX 4060 GPU and an Intel i7-13700K CPU (Fig. 3d), using the same number of iterations for both setups. With 37 iterations, patch deconvolution achieved runtimes of 7 ms for the 19 MLA setup and 8 ms for the 37 MLA setup. Reducing the iteration to 19 further halves the reconstruction time to approximately 4 ms in 19 MLA setup, equivalent to 250 reconstructions per second. A library of cell reconstructions is presented in Supplementary Note S4.B.

### 4.2. Size and order of patches

Next, we evaluated two key parameters in patch deconvolution using simulated data with known ground truth: (1) the order in which the patches are processed and (2) the size of the patch, to assess their impact on reconstruction quality. A square MLA layout was adopted for a simpler patch segmentation. Simulations used 72 confocal-imaged T-cells with Poisson noise added to achieve varying PSNR levels from 1dB to 80dB in the FLFM images (Fig. 4a).

**Fig. 4.**
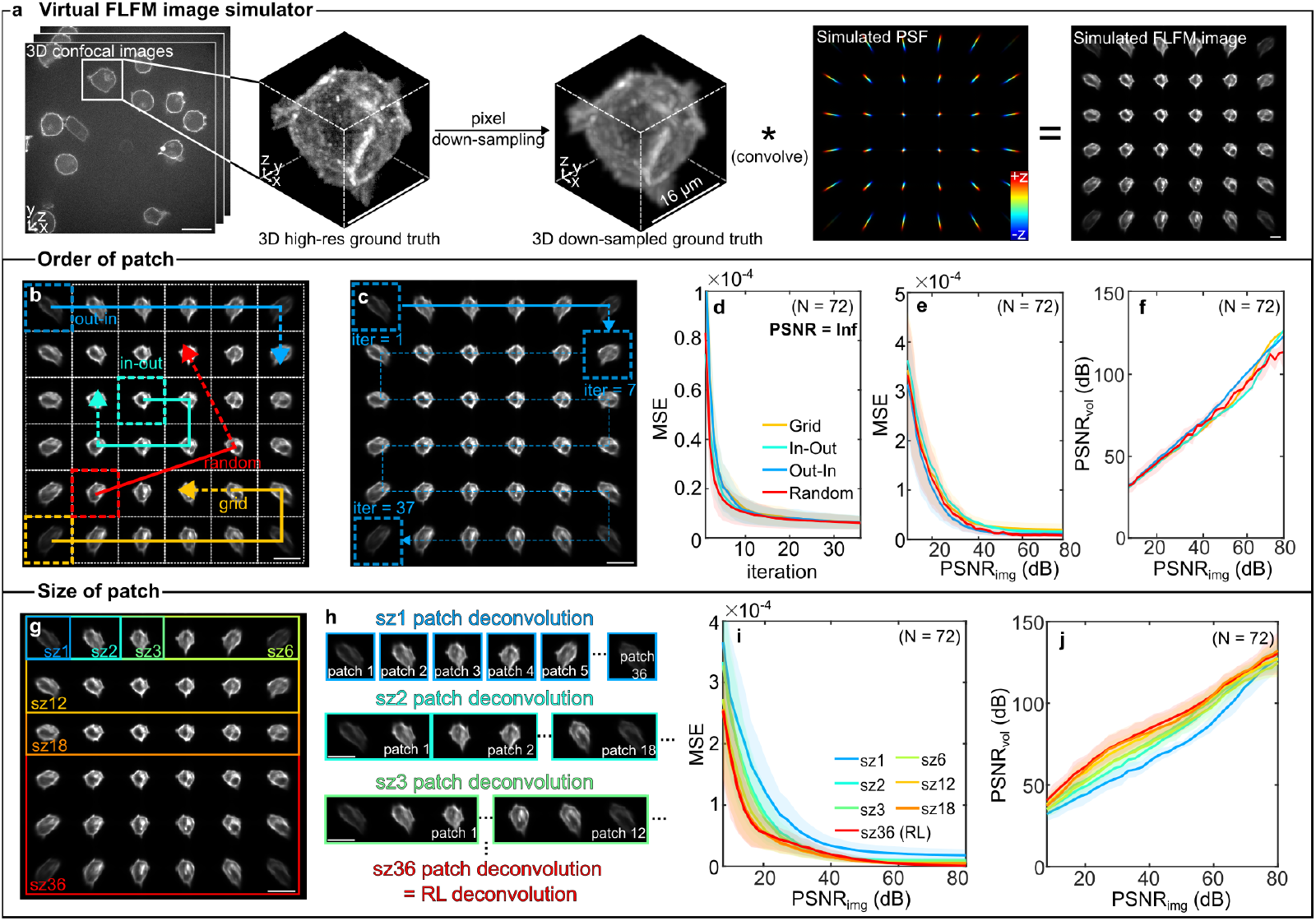
Size and order of patches. **a** Workflow of the virtual FLFM simulator. high-resolution 3D confocal images are down-sampled and convolved with simulated FLFM PSFs to generate test data. **b** Different order of patch deconvolution: *grid, in-out, out-in*, and *random* orders. **c** Visualisation of the *out-in* deconvolution process. **d** Convergence comparison using different patch orders under a noise-free condition. **e, i** MSE between ground truth and patch deconvolution results at final iteration across different peak signal-to-noise ratios (PSNR). **f, j** PSNR of reconstructed volumes at final iteration versus PSNR of the simulated FLFM image. **g** Deconvolution using various patch sizes (*sz1* to *sz36*); *sz36* represents full-image (RL) deconvolution. **h** Visualisation of patch deconvolution across patch sizes. All scale bars are 15 µm.

Four patch processing orders were tested (Fig. 4b): *out-in* (spiralling from the outermost views inward), *in-out* (the reverse), *grid* (raster scan), and *random*. Each patch was used once, so the number of iterations equalled the number of patches *p* (Fig. 4c). Fig. 4e shows that *random* converging slightly faster than the others, but consideration of the entire PSF shows no difference. This is also shown in similar reconstruction quality across PSNR levels from 1 dB to 80 dB in Fig. 4e. Fig. 4f shows that the reconstructed PSNR was always higher compared to the input FLFM image due to multi-view fusion through deconvolution, but independent of patch order. Therefore, if an early stopping is needed for a faster reconstruction, the *random* order can be used.

The size of patch is another important factor. Up until this point, we have considered patch sizes containing only one FLFM perspective view (*sz1* in Fig. 4g). However, patches may include multiple perspectives, such as *sz2* (2 views), *sz3* (3 views), up to *sz36*, representing full-image RL deconvolution (Fig. 4h). We tested different patch sizes over 36 iterations (equal to the number of views) using a *random* order. MSE comparisons across PSNRs (Fig. 4i) revealed that *sz36* was most robust to noise, while *sz1* was least. The largest improvement occurred between *sz1* and *sz2*. This trend is also shown in PSNR results (Fig. 4j). Thus, *sz2* offers a practical trade-off between speed and reconstruction quality.

### 4.3. Validation on flow cell data

Next, we evaluated the performance of patch deconvolution under flow conditions. Membrane-labelled T-cells were imaged at 300 events/s (0.5 ms exposure, 16-bit mode, 1000× 1000 image) (Fig. 5a) and 1500 events/s (0.1 ms exposure, 8-bit mode, 1000× 1000 image). A PSF library was simulated for implementing hybrid PSF, and each frame was assigned a specific PSF (Supplementary Note S5). At 0.5 ms exposure, the FLFM image PSNR ranged from 18–22 dB, slightly lower than typical short-exposure microscopy [44] due to signal division across views. Despite this, patch deconvolution closely matched RL deconvolution in reconstruction accuracy (Fig. 5c). Even at 0.1 ms, 3D reconstructions remained largely consistent, though patch deconvolution exhibited higher background noise (Supplementary Note S4.C). Besides the morphology comparison, reconstructed cell volumes (Fig. 5d), filling the binary masks for reconstructed cells, showed comparable trends between both methods. Combined with PSNR analysis (Fig. 5e), the slightly reduced image quality of patch deconvolution did not compromise the accuracy of quantitative analysis, thereby enabling sorting performance comparable to RL reconstruction. A library of reconstructed cells in flow is shown in Supplementary Note S4.C.

**Fig. 5.**
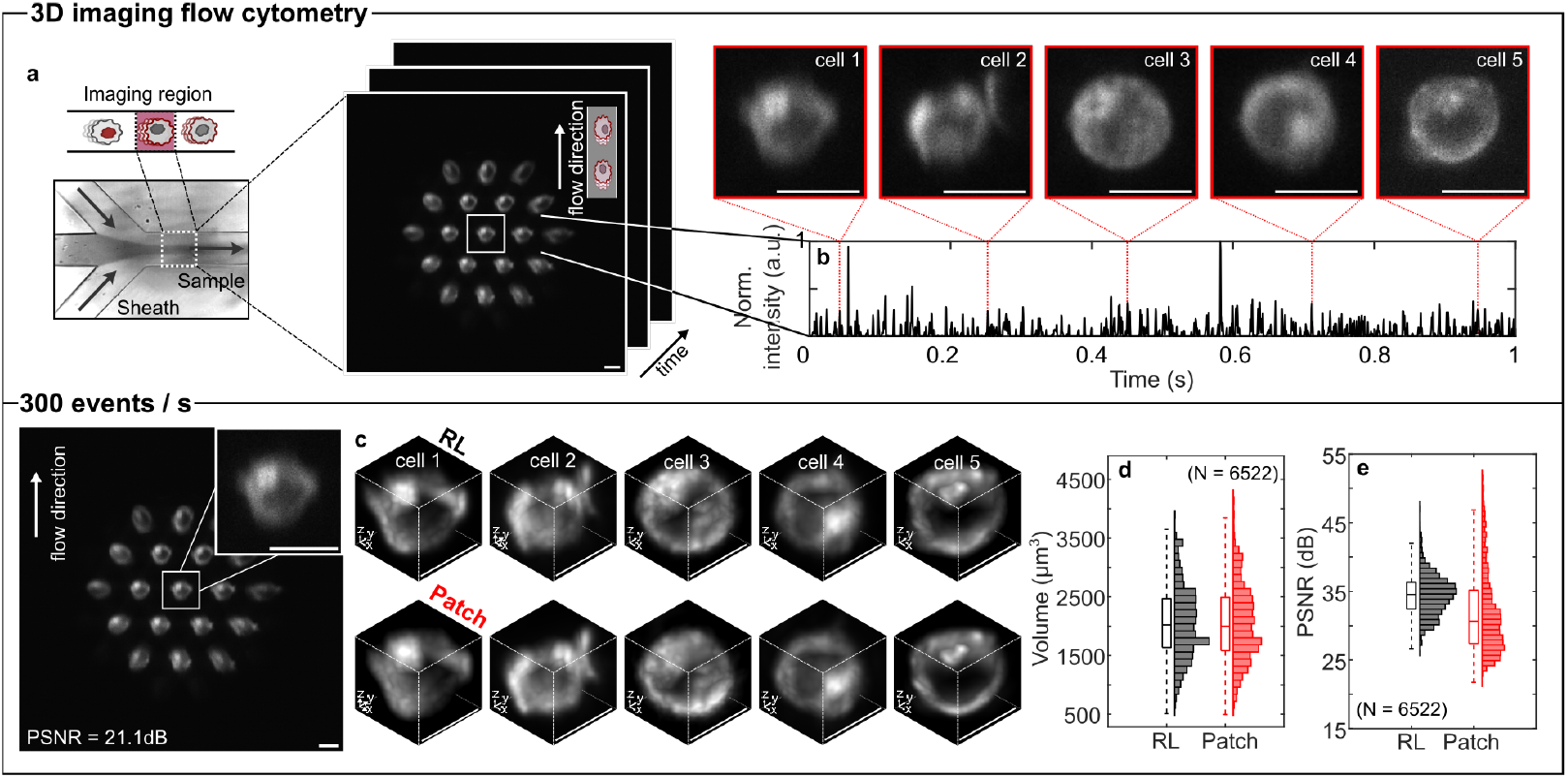
Validation on flow cell data. **a** Schematic of 3D light-field flow cytometry. **b** Sum of normalised intensity in the central view over one second at 0.5 ms exposure, highlighting five example cells. **c** 3D volume renderings of five cells reconstructed using patch and RL deconvolution at 0.5 ms exposure. **d** Comparison of cell volumes from patch and RL deconvolution. **e** PSNR comparison of reconstructed volumes between patch and RL deconvolution. All scale bars are 15 µm.

## 5. Conclusion and Discussion

Here, we have introduced and validated patch deconvolution as a computationally efficient algorithm for 3D cell reconstruction in FLFM at a rate of 100-200 reconstructions per second. Inspired by OSEM [38] and efficient Bayesian multiview deconvolution [37], patch deconvolution speeds up reconstruction by a factor of approximately *p*, where *p* denotes the number of views in the FLFM image. A comparative summary of performance across algorithms discussed in the introduction is provided in Table 1. This throughput enables millions of cells to be imaged and timely reconstructed in a single experiment, overcoming a major barrier to the widespread adoption of 3D imaging flow cytometry in life science.

**Table 1.** Comparison of different reconstruction methods. The PSF of a FLFM is composed of PSFs from multiple microlenses, analogous in concept to joint RL deconvolution. Therefore, in this table, RL deconvolution applied to FLFM is referred to as joint RL deconvolution. ^1^Here, one iteration denotes a full update using the complete PSF, different to iterations defined previously in the text.

| Method | Model | Generalisability | Convergence speed | Hardware requirement | Ref |
| --- | --- | --- | --- | --- | --- |
| Subset RL with plane-wise optimisation | This work | High | $\leq 1$ iteration<sup>1</sup> | CPU/GPU | - |
| Joint RL | Hua *et al.*, Ingaramo *et al.* | High | 20-40 iterations | CPU/GPU | [20, 45] |
| Joint RL with Wiener-Butterworth filter | Guo *et al.* | High | 1-5 iteration | CPU/GPU | [28] |
| Joint RL with projection-estimation | Wu *et al.* | High | 1-5 iteration | CPU/GPU | [29] |
| Subset RL | Preibisch *et al.* | High | $\leq 1$ iteration<sup>1</sup> | CPU/GPU | [37] |
| VCD / F-VCD-Net | Wang *et al.*, Yi *et al.* | Low | N/A | GPU | [30, 31] |
| HyLFM-Net | Wagner *et al.* | Low | N/A | GPU | [32] |
| SeReNet | Lu *et al.* | High | N/A | GPU | [33] |

While patch deconvolution shows great promise for high throughout 3D imaging cytometry, there are three key limitations should be considered: (1). physical limitations on reconstruction time, (2) limited FOV size for having spatial isolation between patches, (3). less robust to high noise conditions. While faster GPUs can also bring RL reconstruction into the order of millisecond, patch deconvolution always provides an intrinsic and consistent *p*-fold speed-up. In practical, however, reconstruction times are also limited by data transfer between RAM and GPU, code efficiency, and overheads from instruction calls. In addition to the implemented-level limitations, patch deconvolution faces challenges in handling overlapping views, where it becomes difficult or impossible to segment distinct patches. This issue can be avoided by placing an iris at the image plane to restrict the FOV, thus eliminating overlap. Moreover, patch deconvolution is less robust to noise, indicated by images under 0.1 ms exposure and 8-bit mode (Supplementary Note 4.C), as each update is based on noisier, less consistent statistics. Improved labelling strategies or low-noise cameras are therefore essential for a higher acquisition speed.

Patch deconvolution enables millisecond-scale 3D reconstructions, achieving cell sorting speeds comparable to the fastest reported methods (Table 2), while relying on a simpler microscope setup and reconstruction algorithm. This significantly increases the competitiveness of light-field cytometry (LFC).

**Table 2.** Comparison of different imaging flow cytometry methods. Comparison of different imaging flow cytometry methods based on their capability for 3D imaging, resolution, detection range, acquisition speed (events per second), reconstruction speed (events per second) and simplicity of setup. The reconstruction speed here is proportional to sorting speed.

| Instrument | 3D | Resolution | Range | Acquisition speed | Reconstruction speed | Simplicity | Ref |
| --- | --- | --- | --- | --- | --- | --- | --- |
| This work (19/37MLA) | ✓ | 1.3 $\mu\text{m}$ | Whole-cell | 1500 | 100-200 | ++++ | - |
| FLFM (3MLA) | ✓ | 600 nm | Subcellular | $\leq 5000$ | 5-10 | ++++ | [20] |
| Radiofrequency-tagged emission | × | 1.55 $\mu\text{m}$ | Whole-cell | 15000 | 15000 | +++ | [7] |
| Light-sheet microscopy | ✓ | 1.8 $\mu\text{m}$ | Whole-cell | 10-20 | 10-20 | +++ | [11] |
| Wide-field illumination with psf engineering | ✓ | 300 nm | Single-molecule | 250 | 250 | ++++ | [15] |
| Lattice light-sheet microscopy | ✓ | 1 $\mu\text{m}$ | Whole-cell | $< 10$ | $< 10$ | ++ | [10] |
| Scanning light-sheet illumination | ✓ | 2 $\mu\text{m}$ | Whole-cell | 500 | 500 | ++ | [12] |
| Light-sheet microscopy on chip | ✓ | 300 nm | Whole-cell | $< 10$ | $< 10$ | ++ | [13] |
| Commercial system (ImageStream) | × | 500 nm | Whole-cell | 2000 | 2000 | ++++ | [46] |

Across both static and flow datasets and under varied noise conditions, patch deconvolution produced reconstructions that were comparable to RL in resolution, morphology, and cell volumes, thus having negligible impact on sorting accuracy. Since the only difference lies in matrix size, patch deconvolution requires no changes to existing hardware or software systems and remains fully compatible with other optimisation techniques. Patch deconvolution is thus a practical and powerful addition to the 3D LFC. With the acceleration ability, it opens new opportunities for cell sorting based on complex spatial features [2], such as protein co-localisation [8] or rare subcellular patterns (Supplementary Note S6), within large populations. These capabilities are vital for applications in flow cytometry where both precision and throughput are critical.

## Acknowledgement

The authors would like to thank Kevin O’Holleran for valuable discussions.

## Author contributions

B.F. and S.F.L. conceived the project. T.P.J.K., L.E.W., R.A.F. and S.F.L. oversaw the project. C.L.J., D.H., C.O’B.-B. and V.J.prepared and stained the cell samples. B.F. and C.L.J. took experimental data. B.F., C.L.J and S.Y. built a Fourier light-field microscope. C.L.J and H.Y. built the flow cytometer. B.F. performed simulations, reconstructions and analysis. B.F. and S.F.L. wrote the manuscript with input from all authors.

## Data availability

Codes and data in support of this study can be found in the following locations:

- FLFM simulation and patch deconvolution code: https://github.com/binfu0728/patch_deconvolution
- Raw Data and code for all Figures: 10.5281/zenodo.14020429

## Supplemental document

See Supplement 1 for supporting content.

